# Integrating Culture-based Antibiotic Resistance Profiles with Whole-genome Sequencing Data for 11,087 Clinical Isolates

**DOI:** 10.1101/463901

**Authors:** Valentina Galata, Cédric C. Laczny, Christina Backes, Georg Hemmrich-Stanisak, Susanne Schmolke, Andre Franke, Eckart Meese, Mathias Herrmann, Lutz von Müller, Achim Plum, Rolf Müller, Cord Stähler, Andreas E. Posch, Andreas Keller

**Affiliations:** Chair for Clinical Bioinformatics, Saarland University, Building E2.1, Saarbrücken 66123, Germany.; Institute of Clinical Molecular Biology, Christian-Albrechts University of Kiel, Schittenhelmstrasse 12, Kiel 24105, Germany.; Ares Genetics GmbH, Karl-Farkas-Gasse 18, Vienna 1030, Austria.; Siemens Healthcare GmbH, Strategy and Innovation, Hartmannstrasse 16, Erlangen 91052, Germany.; Department of Human Genetics, Saarland University, Building 60, Homburg/Saar 66421, Germany.; Institute of Medical Microbiology and Hygiene, Saarland University, Kirrberger Strasse, Campus Building 43, Homburg/Saar 66421, Germany.; Curetis GmbH, Max-Eyth-Strasse 42, Holzgerlingen 71088, Germany.; Department of Pharmacy, Pharmaceutical Biotechnology, Saarland University, Saarbrücken 66123, Germany; Department of Microbial Natural Products, Helmholtz-Institute for Pharmaceutical Research Saarland (HIPS), Saarland University, Building E8.1, Saarbrücken 66123, Germany; Helmholtz Center for Infection Research and Pharmaceutical Biotechnology (HZI), Saarland University, Saarbrücken 66123, Germany.

**Keywords:** Antibiotic resistance, Whole-genome sequencing, Bacteria, Pan-genome

## Abstract

Emerging antibiotic resistance is a major global health threat. The analysis of nucleic acid sequences linked to susceptibility phenotypes facilitates the study of genetic antibiotic resistance determinants to inform molecular diagnostics and drug development. We collected genetic data (11,087 newly sequenced whole genomes) and culture-based resistance profiles (10,991 of 11,087 isolates were comprehensively tested against 22 antibiotics in total) of clinical isolates including 18 main species spanning a time period of 30 years. Species and drug specific resistance patterns could be observed including increasing resistance rates for *Acinetobacter baumannii* to carbapenems and for *Escherichia coli* to fluoroquinolones. Species-level pan-genomes were constructed to reflect the genetic repertoire of the respective species such as conserved essential genes and known resistance factors. Integrating phenotypes and genotypes through species-level pan-genomes allowed to infer gene-drug resistance associations using statistical testing. The isolate collection and the analysis results have been integrated into a resource, GEAR-base, available for academic research use free of charge at https://gear-base.com.

## Introduction

The development of new antimicrobial drugs has largely stagnated over the last few decades[1], while the drug resistance rates of many pathogens have at the same time been increasing[2–4]. Various large-scale efforts exist investigating emerging drug resistance, such as the Meropenem Yearly Susceptibility Test Information Collection (MYSTIC) program[2], the Canadian National Intensive Care Unit (CAN-ICU) study[5], the Canadian National Surveillance (CANWARD) study[6,7], the Center for Disease Dynamics, Economics and Policy (CDDEP) study[3], and the European Antimicrobial Resistance Surveillance Network (EARS-Net) survey[8]. The results of these studies have shed light on the most common bacterial pathogens and resistance rates for regularly administered antibiotics, with the primary focus on the trend analysis of specific bacterial groups, periods of time, or locations[2,3,9–12]. The global challenge of emerging drug resistance is further exacerbated by the rising prevalence of microorganisms with multidrug resistance (MDR) phenotypes[13]. Accordingly, identifying and administering the most effective drug in each individual case is of even greater importance for successful treatment of bacterial infections. However, these studies did not investigate the genetic repertoire of the pathogens, which represents an important source of information — e.g., the resistance genotype may be readily revealed while the respective phenotype is misleading or not expressed under artificial laboratory conditions[14,15].

Simultaneously, the recovery of genomic information from microorganisms via high-throughput sequencing approaches has become a routine task. This is not only allowing high-resolution study of individual organisms’ genomes, but also aggregated study in the form of “pan-genomes” — the united genetic repertoire of a clade[16]. Pan-genomes can be used to identify common genetic potential — i.e., the “core” genes of a clade — as well as genes that are less broadly conserved (“accessory” or “singleton” genes)[16]. This facilitates the identification of essential genes or genes that provide adaptation advantages. Multiple computational approaches exist for the systematic creation of pan-genomes — e.g., Roary[17], EDGAR[18], panX[19] — and a variety of bacterial pan-genomes, typically at the species-level, have thus far been constructed[20–23]. Most pan-genome studies focus on distinct species and do not always cover clinically relevant species. For example, MetaRef represents a resource that provides information about pan-genomes from multiple species and integrates approximately 2,800 public genomes[24]. Although the diversity of the therein included organisms is particularly broad, the depth is limited in relation to clinically relevant bacteria — e.g., seven *Klebsiella pneumoniae* genomes. Moreover, individual isolates included in the studies often span narrow time frames and/or have limited geographic spread.

While pan-genomic studies typically focus on the genetic information alone, efforts combining genomic and phenotypic information, in particular from antibiotic resistance testing, for the study of conserved or emerging resistance mechanisms are becoming increasingly prevalent[25–28]. There are many antibiotic resistance resources available[29] however only few link genomic and phenotypic information of bacterial isolates. One of such resources is the Pathosystems Resource Integration Center (PATRIC)[30] which represents a rich service for the study of > 80,000 genomes[31]. Yet, antimicrobial resistance information is available only for about 10% of the genomes. Furthermore, as PATRIC’s genomes and metadata are imported from public resources which are populated by individual research efforts, standardization or normalization is challenging. Finally, individual taxa may be underrepresented and, thus, warrant expansion — e.g., the number of *Escherichia spp.* genomes with antimicrobial resistance metadata is almost two orders of magnitude smaller than the respective number of *Mycobacterium spp.* genomes[31].

Motivated by the importance of linking resistance phenotypes and genomic features, we collected whole genome sequencing data of 11,087 clinical isolates representing, *inter alia*, 18 main bacterial species. The samples were collected in North America, Europe, Japan, and Australia over a period of 30 years, and processed in a concerted effort, thereby reducing experimental bias. Culture-based resistance testing was performed for 10,991 of the 11,087 isolates including 22 antibiotic drugs. Furthermore, species-level pan-genomes were constructed on the basis of per-isolate *de novo* assemblies and were used to infer gene-drug resistance associations. This wealth of information is integrated into an online resource, Genetic Antibiotic Resistance resource, or in short, GEAR-base (**Figure 1**). Providing broad organismal, antibiotic treatment, and temporal coverage, the present resource is expected to support the pan-genome-based study of bacteria and to advance research on known or emerging antibiotic resistance mechanisms. GEAR-base is available for academic research use free of charge at https://gear-base.com.

**Figure 1.**
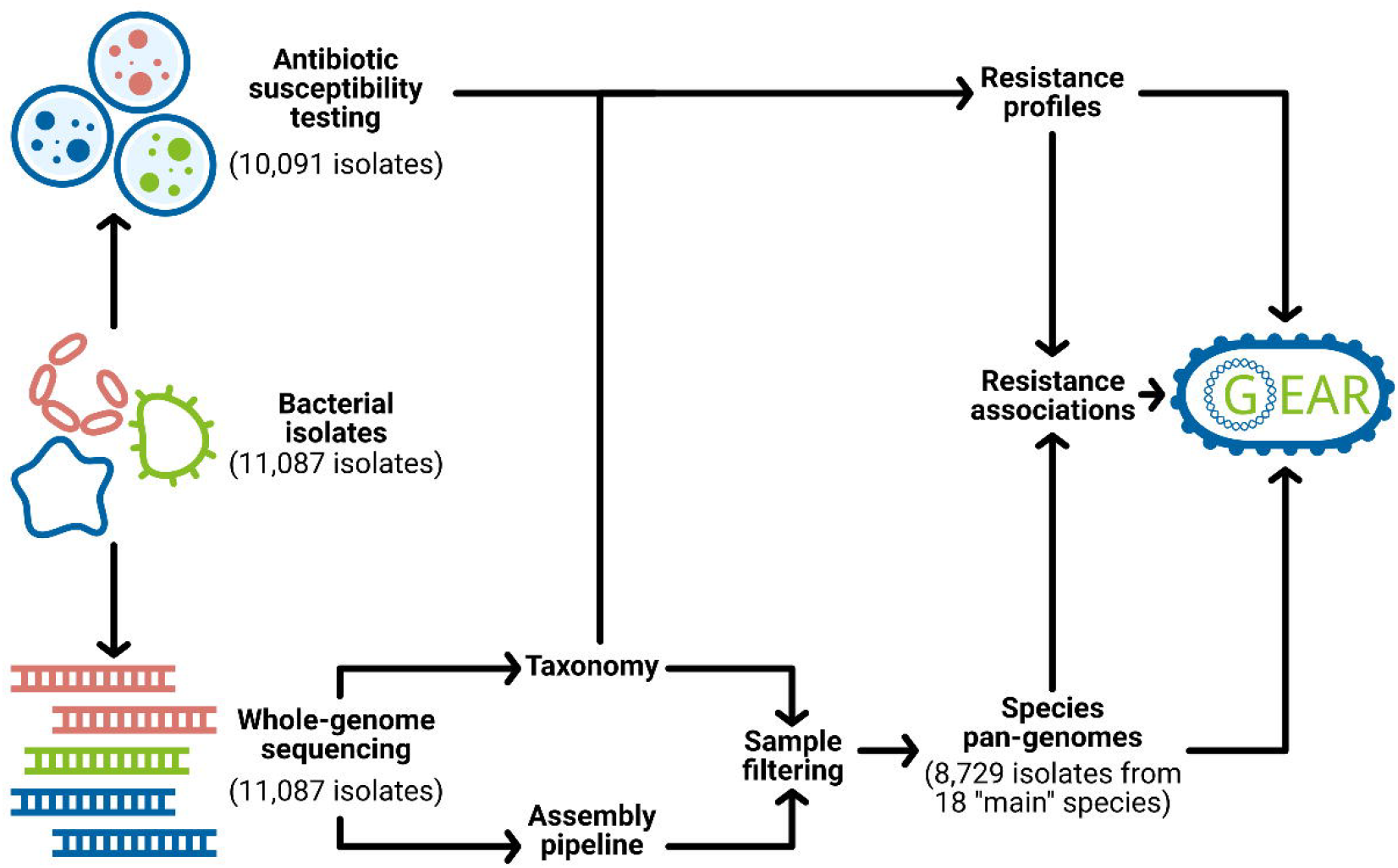
GEAR-base workflow and structure. Schematic overview of data collection, processing and integration into GEAR-base.

## Results

### Resistance testing of cultured bacterial isolates

The present dataset of 11,087 bacterial isolates covered a total of 6 families, 14 genera and 20 species (considering species with at least 50 isolates, **Supplementary Table S1**) and comprised two data sets: 1,001 isolates from the *Staphylococcus aureus* strain collection and 10,086 isolates from the Gram-negative collection. From the *S*. *aureus* strain collection, 993 isolates were tested for methicillin resistance and susceptibility (MRSA/MSSA; Methods). For 9,998 isolates from the Gram-negative collection, culture-based antimicrobial susceptibility testing (AST) for 21 commonly prescribed Food and Drug Administration approved antibiotics from 8 drug classes was performed to determine the respective minimum inhibitory concentrations (MICs) (**Figure 2 A**). The resistance profiles were determined for each isolate in accordance with the EUCAST guidelines (v. 4.0) for a total of 182 drug concentrations (from 7 to 11 concentrations per drug; **Supplementary Table S2** and **S3**, **Figure 2 B**). Whole-genome sequencing (WGS)-based taxonomic identification was performed for all isolates[32]. In the following, we focus on the analysis results of the MIC and resistance profiles of the 9,998 isolates from the Gram-negative collection.

**Figure 2.**
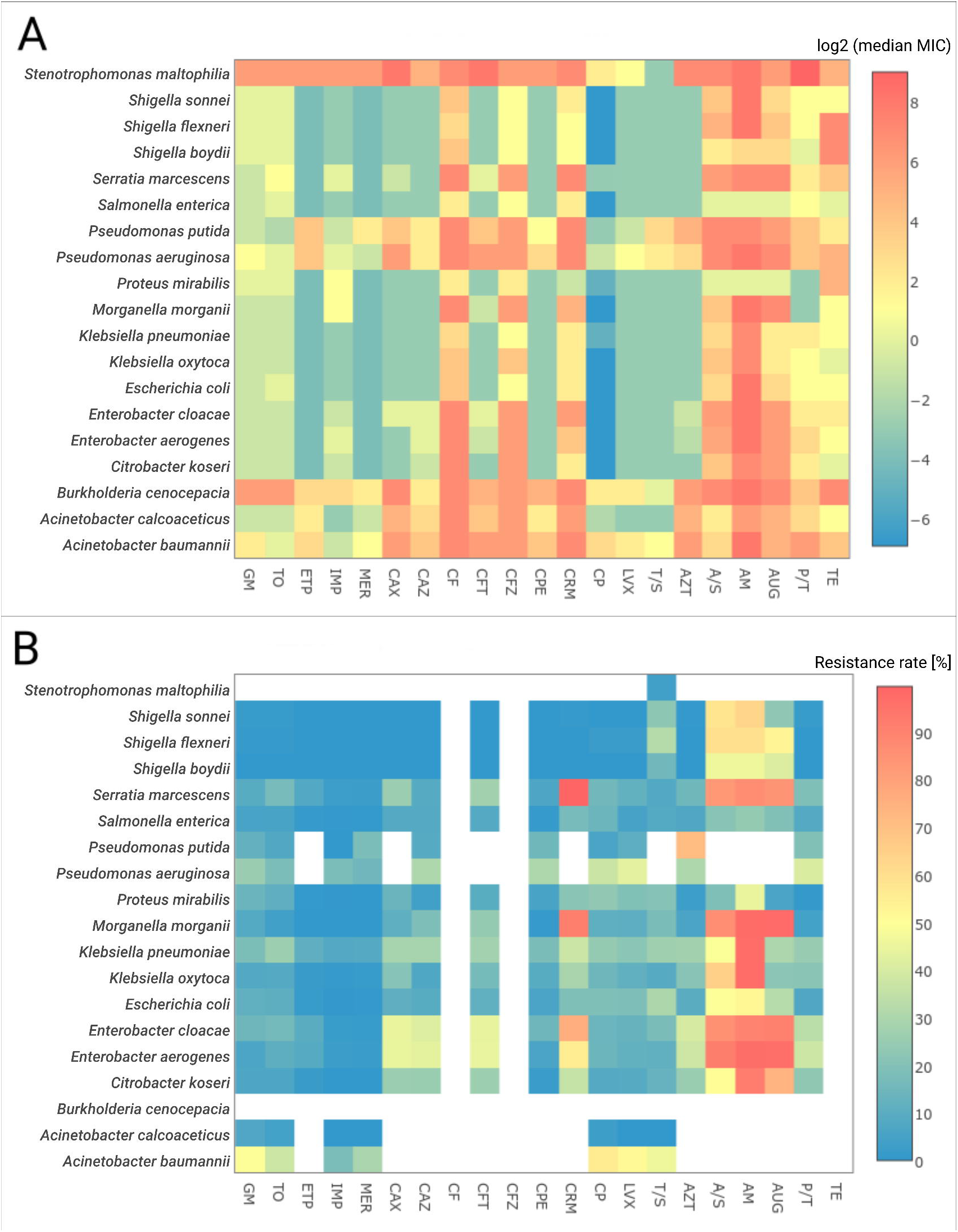
Resistance profiles overview. Heatmap of (**A**) log-transformed (base 2) median MIC values and (**B**) resistance rates for all species with at least 50 isolates. Drugs labels were grouped relative to their class. The cells are colored according to the color key on the right; white color corresponds to the cases where no breakpoints were available from the used guidelines.

All patient-derived isolates were collected in clinics located in North America, Europe, Japan, and Australia from 1983 to 2013 (**Supplementary Figure S1**). Varying degrees of resistance were observed among the isolates (**Figure 2 B**). The majority of species demonstrated relatively low resistance rates (<20%) to aminoglycosides (gentamicin and tobramycin) and carbapenems (ertapenem, imipenem, and meropenem) except for *Acinetobacter baumannii* (≥29% for aminoglycosides and meropenem), *Pseudomonas aeruginosa* (26% for gentamicin), and *Klebsiella pneumoniae* (26% for tobramycin). These rates were compared against two independent large-scale studies — the Center for Disease Dynamics, Economics and Policy (CDDEP, U.S.-based results; CDDEP ResistanceMap, https://resistancemap.cddep.org/AntibioticResistance.php, accessed September 26, 2017)[3] program and the MYSTIC program[2] — for matching species and drug data. Both studies report low (<20%) resistance rates for the aminoglycosides and carbapenems during observed time intervals (1999-2012/2014 for CDDEP and 1999-2007/2008 for MYSTIC) except for *A*. *baumannii* (CDDEP: >20% since 2005 for carbapenems, >35% in 1999-2012 for aminoglycosides; MYSTIC: >37% in 2007/2008 for carbapenems, >20% in most years for aminoglycosides). For *K*. *pneumoniae* and tobramycin (aminoglycosides for CDDEP), MYSTIC and CDDEP reported >10% resistance rates since 2005 with only one value of above 20% observed by MYSTIC in 2007. Finally, for *P*. *aeruginosa* and gentamicin MYSTIC reported a resistance rate of only around 10%. The rate of isolates resistant to multiple antibiotic drugs — i.e., resistant to at least three drugs from different drug classes (CDDEP ResistanceMap, https://resistancemap.cddep.org/AntibioticResistance.php, accessed September 26, 2017) — was highest for *A. baumannii* (44%) and for *Enterobacter* spp. (41–45%). For the remaining species and drug classes, the multiple drug resistance (MDR) rates were at least 20%, except for *Acinetobacter calcoaceticus* (0%), *Salmonella enterica* (11%), and *Shigella* spp. (0–3%). In addition to the investigation of individual species–drug combinations, we analyzed whether drug pairs showed correlating MIC profiles over all isolates (**Supplementary Figure S2**). In general, the highest correlations were expectedly found within single drug classes — e.g., for fluoroquinolones, aminoglycosides, and carbapenems. While for some species — e.g., *Burkholderia cenocepacia* — a clear clustering according to drug classes and their mechanism of action was observed, other species, such as *S*. *enterica*, showed less pronounced cluster structures.

Subsequently, we compared resistant and non-resistant isolates with respect to their collection year in order to identify potential trends of de-/increasing antibiotic resistance rates (**Supplementary Figure S3**, **Supplementary Figure S4**, and **Supplementary Table S4**). The following species-drug pairs were found to exhibit particularly low *P* values (WMW-test FDR adjusted *P* value < 1e-17) as well as increases in resistance over time: *K*. *pneumoniae* to cefepime, *K*. *pneumoniae* and *A*. *baumannii* to carbapenems, and *Escherichia coli* to fluoroquinolones. Similar trends were reported by the CDDEP[3] program (CDDEP ResistanceMap, https://resistancemap.cddep.org/AntibioticResistance.php, accessed September 26, 2017) and the MYSTIC program[2], including increasing resistance rates for *A*. *baumannii* to carbapenems (43% from 1999 to 2014 in the US, CDEEP), and for *E*. *coli* to fluoroquinolones (30% from 1999 to 2014 in the US, CDEEP; >20% from 1999 to 2008, MYSTIC).

While the culture-based results provide species-resolved information about resistance rates over time and corroborate previous findings on the global increase in antibiotic resistance, genetic features represent important factors and were thus concomitantly considered.

### Whole-genome de novo assembly of isolates and species pan-genomes

A total of 11,087 bacterial isolates were whole-genome sequenced using Illumina Hiseq2000/2500 sequencers, resulting in a median number of 1,517,147 paired reads per isolate (stdev 620,481). *De novo* assemblies were successfully created for 11,062 (99.8%) isolates (**Figure 3**) and of these, 10,764 (97.3%) passed the stringent assembly quality criteria. Moreover, 9,206 (83% of 11,087) of the isolates fulfilled the quality criteria for taxonomic assignment. A total of 8,729 isolates, representing 18 main species with at least 50 isolates, were used after stringent quality filtering (see **Methods** for sample filtering details) in the subsequent analyses and in the construction of species-level pan-genomes, with *Acinetobacter calcoaceticus* and *Pseudomonas putida* having less than 50 isolates passing (**Supplementary Table S3**).

**Figure 3.**
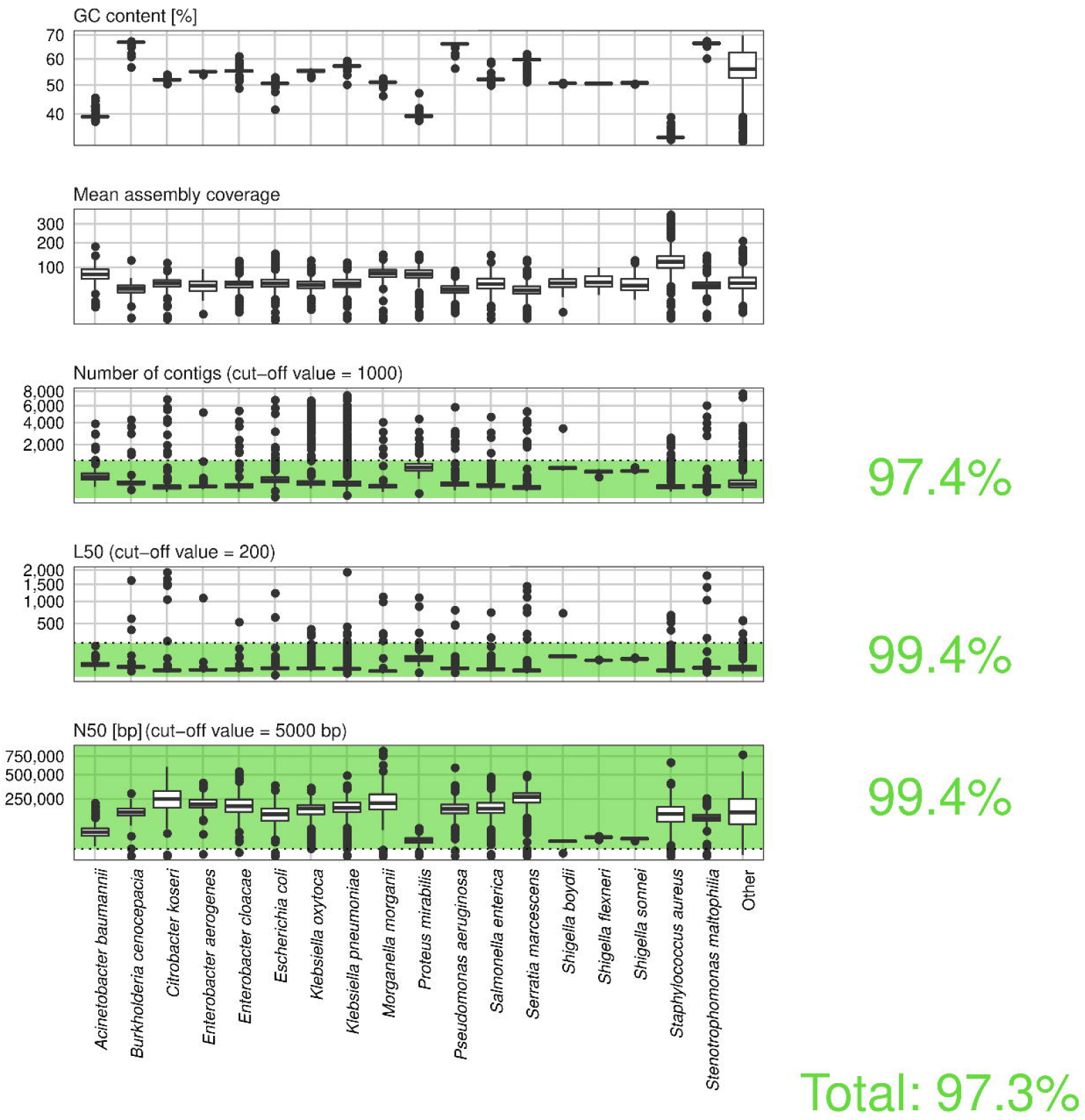
Assembly quality overview. Assembly summary statistics for the 11,062 isolates with a de novo assembly. The isolates were grouped by their species taxon; all isolates not belonging to any of the main 18 species used for pan-genome construction were grouped into “Other”. The box plots show the GC content, mean assembly coverage, number of contigs, L50 value, and N50 value, for contigs of at least 200bp. The assembly quality cut-off values are illustrated by dotted lines. The plot area satisfying the respective filtering criterion is colored in green. Percentages of isolates passing the individual as well as all criteria are shown to the right.

First, the presence/frequency of genes from a set of 111 single-copy marker genes (“essential genes”), according to Dupont *et al.*[33], was used as a proxy to estimate the genome completeness of individual *de novo* assemblies. Overall, the assemblies were found to be largely complete: 92 essential genes (82.9%) were identified in at least 99% of the 8,729 isolates (**Supplementary Figure S5**) which were used to construct a phylogenetic tree of these isolates (**Supplementary Figure S6**). Furthermore, species-specific presence and absence patterns were frequently observed (**Supplementary Figure S7 A**). For example, TIGR00389 (glycine–tRNA ligase) was only found in *S. aureus*, whereas TIGR00388 (glycine–tRNA ligase, alpha subunit) was not present in this species. Four genes, TIGR00408 (proline–tRNA ligase), TIGR02387 (DNA-directed RNA polymerase, gamma subunit), TIGR00471 (phenylalanine–tRNA ligase, beta subunit), and TIGR00775 (Na+/H+ antiporter, NhaD family), were not found in any of the isolates, except for sporadic hits in *Pseudomonas aeruginosa* for TIGR00408.

In the next step, ResFams core-based resistance factors[34] were annotated in the isolate assemblies in order to study the species-level distribution of these genetic features. The number of covered Resfams (mean count of hits ≥ 1) varied between species from 4.1% (5 of 123 Resfams, *Morganella morganii*) to 11.4% (14 of 123 Resfams, *A*. *baumannii* and *Shigella sonnei*) (**Supplementary Figure 8**). Three Resfams were found in at least 90% of all considered isolates – RF0007 (ABC antibiotic efflux pump), RF0107 (ABC antibiotic efflux pump), and RF0115 (RND antibiotic efflux pump), the latter with a mean count of hits of at least 5 for 14 of 18 species.

The MLST analysis revealed, that in all species with a typing scheme included in the used version of PubMLST isolates were assigned to at least 6 different sequence types (STs), except for *S*. *sonnei*, and new STs could be identified, except for *Shigella flexneri* and *S*. *sonnei* (**Supplementary Figure S9**). Among these species, the proportion of isolates without a confident assignment was high (≥ 10%) for *B*. *cenocepacia*, *Enterobacter cloacae*, *Klebsiella oxytoca* and *Stenotrophomonas maltophilia*.

The size of the species pan-genomes (i.e., the number of centroids) ranged from 5,838 (*S*. *aureus*, total pan-genome length 5 Mb) to 42,046 (*E. cloacae*, total pan-genome length 30 Mb) (**Supplementary Figure S10**). A centroid refers here to the representative gene of a homologous gene cluster with ≥ 90% pair-wise amino acid sequence identity (Methods). Most centroids were found in fewer than 10% or in at least 90% of the isolates (**Figure 4**). Moreover, all pan-genomes were found to be open based on the analysis of the number of centroids in relation to the number of included genomes (**Supplementary Figure S11**, **Supplementary Table 5**). The two-dimensional embedding of the core centroids from the pan-genomes revealed many taxon specific patterns (**Supplementary Figure S12**) with distinct clusters for *B*. *cenocepacia*, *M*. *morganii*, *A*. *baumannii*, *Proteus mirabilis*, *S*. *aureus*, *S*. *maltophilia*, *P*. *aeruginosa*, and *Serratia marcescens*. We compared the number of (core) centroids in our pan-genomes to the numbers reported by panX[19] (website accessed on January 29^th^, 2018). The number of centroids present in at least 90% of the analyzed genomes was consistent for all matching species (**Supplementary Table 6**). However, the pan-genome size, i.e. the total number of centroids described in GEAR-Base, was similar for *E*. *coli* and *S*. *aureus*, but exceeded substantially the number of centroids described in panX for *A*. *baumannii*, *K*. *pneumoniae*, *P*. *aeruginosa* and *S*. *enterica* (**Supplementary Table 6**). With respect to the presence of essential genes in the species-level pan-genomes, the mean number of centroids containing at least one matching gene was one — i.e., these essential genes were mostly found in only one centroid cluster (**Supplementary Figure S7 B**). However, the mean number of centroids was greater than or equal to 1.25 for eight essential genes — i.e., in some species these genes were found in multiple centroid clusters.

**Figure 4.**
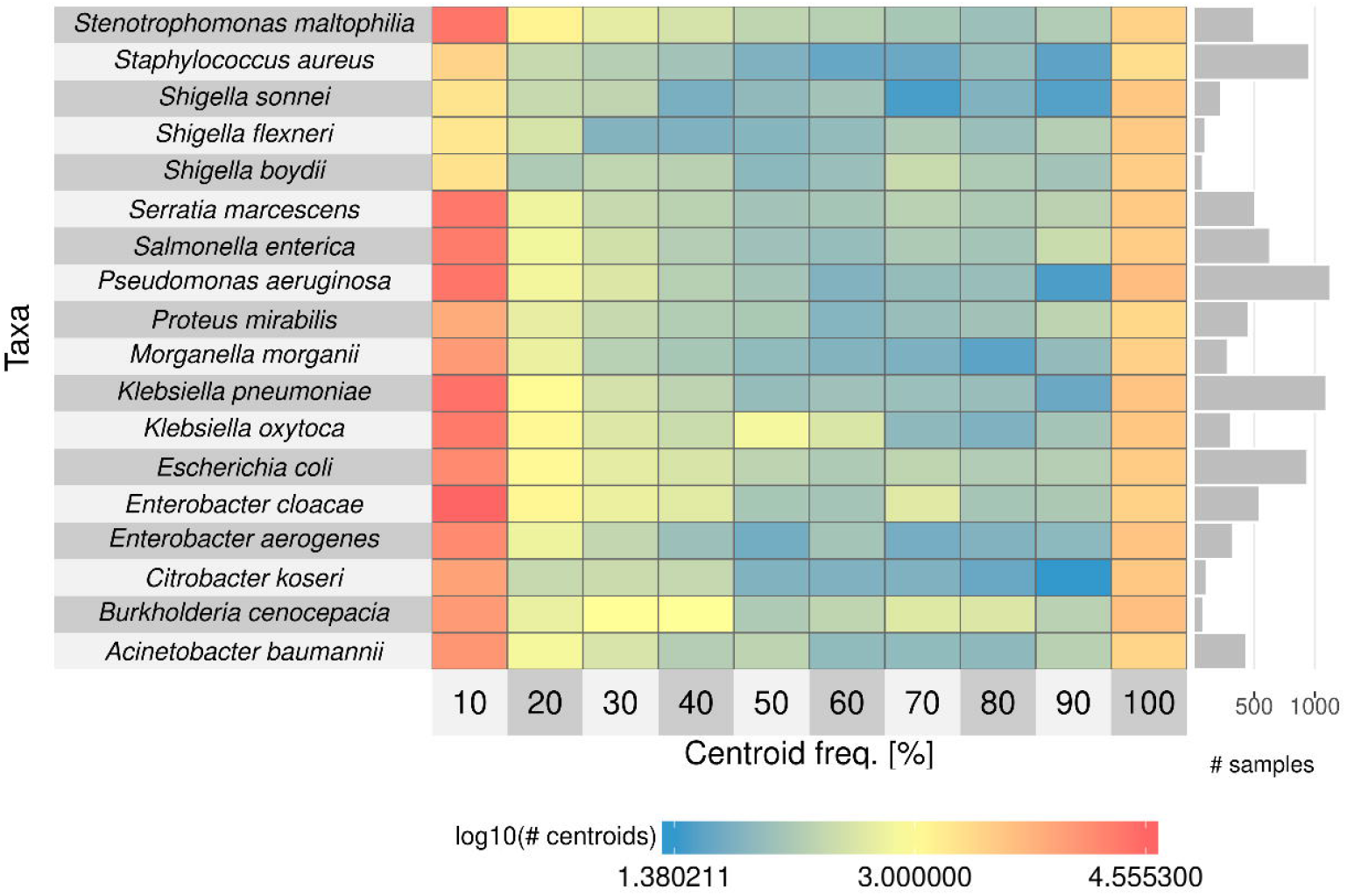
Centroid frequency. Number of centroids in each pan-genome of the 18 main species in relation to their frequency – i.e., the first column contains centroids that are present in fewer than 10% of the isolates, and the last one contains centroids that are present in at least 90% of the isolates. The color of the cells corresponds to the log10-transformed number of centroids. The bar plot on the right side shows the number of samples used to construct the respective pan-genomes.

In the following, the resistance phenotypes and genomic features were linked and significantly associated centroids were further studied with respect to their overlap to known resistance genes from the Resfams core database.

### Resistance associations by linking phenotype and genotype

We used binary information in the form of centroid presence/absence to test for significant centroid-drug associations per species. The number of found associations ranged from below 10 to above 500; most associations (at least 500) were found for *P*. *aeruginosa* and tobramycin, and *K*. *pneumoniae* and gentamicin (**Figure 5**). Furthermore, the drug resistance-associated centroids encoding for a resistance gene were investigated. From the Resfams’ core database 45 of the 123 factors were found in at least one centroid (**Supplementary Figure S13**). Among these, the top ten Resfams genes from both analyses covered various resistance mechanism classes – nucleotidyltransferases, phosphotransferases, acetyltransferases, beta-lactamases, and MFS transporters (**Supplementary Figure S13 B**).

**Figure 5.**
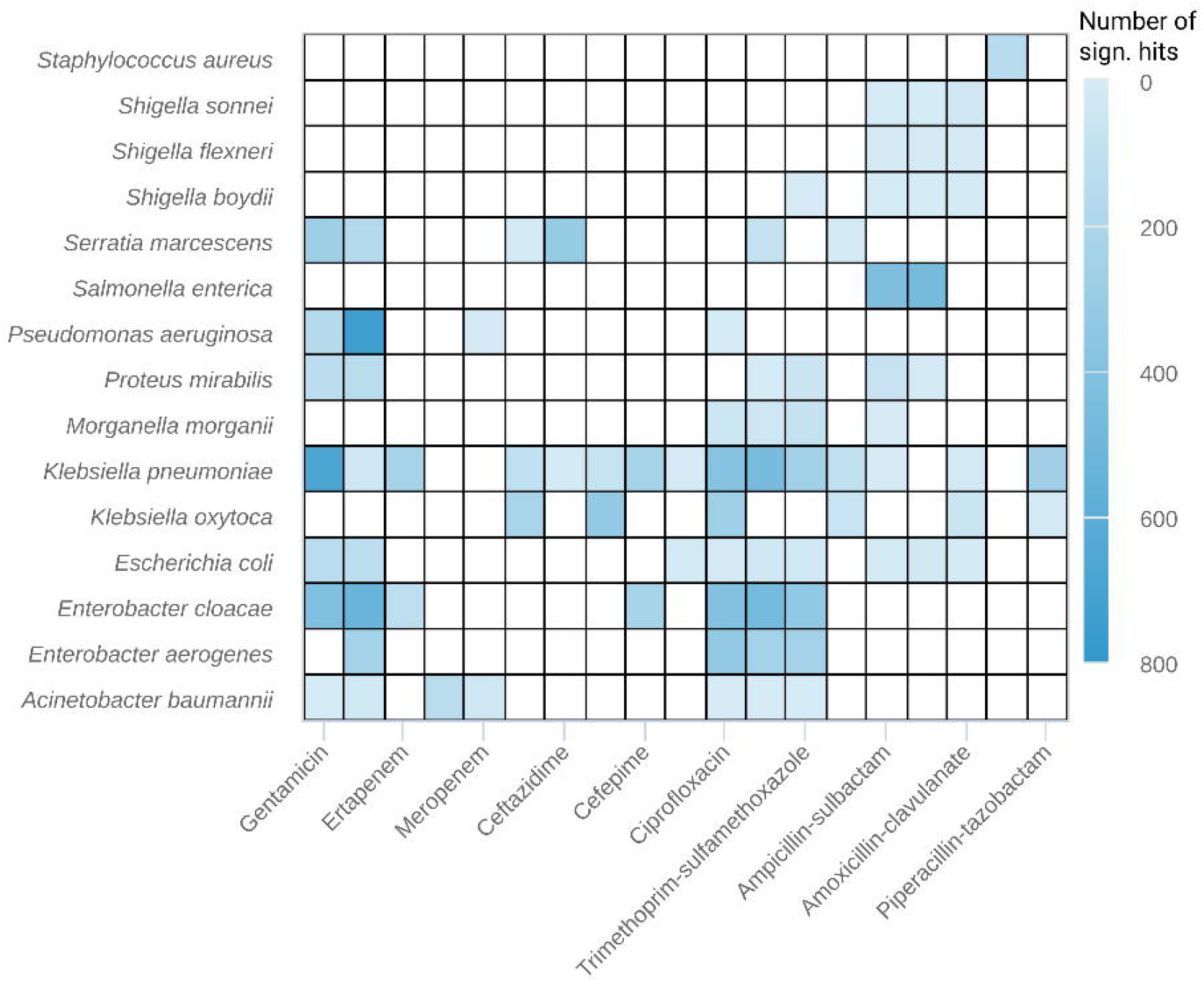
Significant results of the resistance association analysis. Significant results of the resistance association analysis based on centroid presence/absence. The heatmap showes the number of significant hits (small numbers — light blue; large numbers — blue) per taxon and drug. Drugs are sorted according to their class.

### GEAR-base online resource

The GEAR-base resource is freely accessible at https://gear-base.com for academic research use and currently provides two modules for browsing of the database — a culture-based module and a pan-genome module — as well as a module for the analysis of user-provided data. The culture-based module is focused on the Gram-negative isolate collection and provides an interactive view of the taxonomic composition, MIC and resistance profiles, as well as additional meta-data, e.g., collection year or sample distributions. The pan-genome module provides general statistics, such as assembly quality of the included isolates, pan-genome size, and resistance association analysis overview, for both, the Gram-negative and the *S. aureus* isolates. Gene nucleotide sequences can be downloaded for each individual pan-genome centroid and a batch-download of all centroid nucleotide sequences is available. Moreover, pan-genome centroids can be browsed online for specific gene products and filtered by their presence in the isolates. In addition, centroid clusters can be viewed including associated gene annotations, the hits to the Resfams core database, and information about potential resistance associations against the set of herein included drugs. GEAR-base’s analysis module allows the user to query individual gene sequences against the pan-genome centroid sequences using Sourmash[35], against hidden Markov models (HMMs) of pan-genome centroid clusters and Resfams core database using HMMER (http://hmmer.org/), and against the NCBI nt/nr database using BLASTp[36]. Furthermore, a genome-scale search against the present clinical isolate collection, the finished genomes from NCBI’s RefSeq database[37], as well as the NCTC 3000 genomes project from Public Health England and the Wellcome Trust Sanger Institute (http://www.sanger.ac.uk/resources/downloads/bacteria/nctc/, accessed October 18, 2017) can be performed online using Mash/MinHash[38].

To demonstrate GEAR-base’s analysis functionalities, we used a recently published *K*. *pneumoniae* genome[39] (strain 1756, NCBI assembly accession ID GCF_001952835.1_ASM195283v1). In a first step, the chromosome and plasmid sequences were uploaded and a perfect match was found to the genome’s NCBI entry, as expected. The next-best matches were to a *K*. *pneumoniae* isolate from the present collection of clinical isolates (828/1000 shared hashes, distance of 4.71e-3), and to a *Klebsiella* sp. genome (ERS706555) from the NCTC 3000 database (709/1000 shared hashes, distance of 8.89e-3). In a second step, all coding DNA sequences (CDS) were searched against GEAR-base’s pan-genome centroids using Sourmash and against the Resfams core database. The majority of the pan-genome hits were related to *K*. *pneumoniae* (6,206 hits of 11,267) followed by *E*. *aerogenes* (1,537 hits), and *K*. *oxytoca* (1,014 hits). *S. aureus*, a Gram-positive species, served as an outgroup and no hits to this species’ pan-genome were found. In total 37 hits to 21 unique Resfams (core database) were found in the query genome CDS with 23 hits on the chromosome and 14 on the plasmid. The top three most occurring Resfams were RF0115 (8 hits, RND antibiotic efflux pump), RF0098 (3 hits, MexE, RND antibiotic efflux), and RF0053 (3 hits, class A beta-lactamase). Furthermore, the CDS of eight antibiotic resistance genes reported in the original genome announcement were investigated. The HMM-based search of pan-genome centroids resulted in two chromosomal CDS,WP_076027158.1 (multidrug efflux RND transporter periplasmic adaptor subunit OqxA) and WP_004146118.1 (FosA family fosmomycin resistance glutathione transferase), being classified as *K*. *pneumoniae*-derived centroids according to their top hits (with respect to the full sequence score), the top hits of the remaining genes (5 plasmid-derived, 1 chromosome-derived) included centroids from other Gram-negative species. However, the centroid cluster annotations matched the expected protein functions for all eight CDS independent of the species. The top three hits for WP_004146118.1 were centroids from *K*. *pneumoniae*, *E*. *aerogenes*, and *K*. *oxytoca* matching the expected annotation and present in almost all isolates (>98%) of the respective pan-genomes. This high prevalence matches the observations made by Ryota *et al*. reporting similarly high frequency (>96%) of *fosA* in these species[40]. For the beta-lactamases WP_004176269.1 (class A broad-spectrum beta-lactamase SHV-11), and WP_000027057.1 (class A broad-spectrum beta-lactamase TEM-1) the top hits in *Klebsiella* were associated with resistance to penicillins and cephalosporins, and for the aminoglycoside transferases WP_000018329.1 (aminoglycoside O-phosphotransferase APH(3’)-Ia), WP_032491824.1 (ANT(3’’)-Ia family aminoglycoside nucleotidyltransferase AadA22), and WP_000557454.1 (aminoglycoside N-acetyltransferase AAC(3)-IId) the top hits in *K*. *pneumoniae* were associated with resistance to aminoglycosides. Moreover, all three chromosome-derived CDS (WP_004176269.1, WP_076027158.1, and WP_004146118.1) matched to centroids found in > 92% of the *K*. *pneumoniae* isolates, two of the five plasmid-derived CDS (WP_032491824.1 and WP_000027057.1) matched to centroids with a frequency of >25%, while the remaining CDS matched to centroids with a frequency of less than 12%.

## Discussion

In order to support the study of antibiotic resistance, we have built GEAR-base, a resource incorporating paired data on resistance phenotypes and genomic features for an extensive, longitudinal collection of clinical isolates from various bacterial species. This concerted effort is expected to reduce experimental bias and the present resource provides a portal for information retrieval as well as data analysis.

Species-level antibiotic resistance phenotypes can be inspected using GEAR-base’s culture-based module. Specifically, resistance rates and trends across multiple species and antibiotic drugs can be assessed on a large scale, which we believe to be important for current and future antibiotic resistance research. Though, some effect of potential sampling bias cannot be excluded, our results on increased resistance rates corroborate previously reported trends. In addition to this phenotypic information, genomic information is included via the pan-genome module. This information can be used independent of the phenotypic information — i.e., purely from a pan-genomic perspective, e.g., for the study of inter- or intra-species gene conservation. The observed number of core centroids was consistent with the statistics reported by panX. However, GEAR-base pan-genomes are based on significantly higher sample numbers and are substantially larger in size, thus giving access to a comprehensive collection of the genome heterogeneity for human bacterial pathogens. In addition, GEAR-base links these two information layers through centroid-drug associations. These associations can subsequently be explored to study resistance mechanisms. Furthermore, individual researchers can compare genes or genomes of interest to the present resource, thereby providing an independent layer of support. This functionality was demonstrated using a recently published carbapenem-resistant *K. pneumoniae* isolate. While the taxonomic classifications of the genome and of a set of chromosome-derived antibiotic resistance genes were consistent with the expected taxonomy of the isolate, the plasmid-derived antibiotic resistance genes exhibited ambiguous taxonomic assignments, not unexpected for plasmid-borne genes. Moreover, the extensive collection of isolates included herein enabled the study of the overall conservation degrees and the time-resolved frequencies of this exemplary antibiotic resistance gene set.

GEAR-base’s analysis functionality covers external genome databases (NCBI’s RefSeq as well as the NCTC 3000 genomes project from Public Health England and the Wellcome Trust Sanger Institute) in addition to the present collection of clinical isolate genomes. However, because the majority of external genomes are not linked to antibiotic resistance information and centroid-drug associations are considered a key component of the present resource, the pan-genome module is restricted to the present isolates. Additionally, GEAR-base’s species-level pan-genome centroids are available for download and provide a great opportunity for subsequent integration with external genomes for further study.

Emerging antibiotic resistance represents a multi-disciplinary and global challenge. We believe that GEAR-base will serve as a valuable resource enabling the detailed analysis of resistance-associated genomic features. GEAR-base includes a comprehensive selection of clinically highly relevant human microbial pathogens and will thus be of great use for the research community.

## Materials and methods

### Bacterial isolates

The dataset of 11,087 isolates consisted of 1,001 isolates from the *S*. *aureus* strain collection of Saarland University Medical Center and a collection of 10,086 Gram-negative bacterial clinical isolates that form part of the microbiology strain collection of Siemens Healthcare Diagnostics (West Sacramento, California, USA)[32]. DNA extraction using the Siemens VERSANT^®^ sample preparation system[41] and whole-genome next-generation sequencing were performed for all isolates as described in Galata *et al*.[32] (2×100 bp paired-end on Illumina Hiseq2000/2500 sequencers).

### Methicillin susceptibility of *S*. *aureus* isolates

For 993 isolates from the *S*. *aureus* strain collection detection of methicillin resistant and susceptible *Staphylococcus aureus* (MRSA/MSSA) isolates was performed. The specimen were plated on CHROMagar MRSA detection biplates (Mast, Germany). All MRSA positive culture isolates were further confirmed using a penicillin binding protein 2a latex agglutination test (Alere, Germany).

### Susceptibility testing and resistance profiles of Gram-negative isolates

For 9,998 isolates from the Gram-negative isolate collection antimicrobial susceptibility testing (AST) was performed. Frozen reference AST panels were prepared following Clinical Laboratory Standards Institute (CLSI) recommendations[42]. The antimicrobial agents included in the panels are provided in **Supplementary Table S2**. Prior to use with clinical isolates, AST panels were tested and considered acceptable for testing with clinical isolates when the QC results met QC ranges described by CLSI[42].

Isolates were cultured on trypticase soy agar with 5% sheep blood (BBL, Cockeysville, Md.) and incubated in ambient air at 35±1°C for 18-24 h. Isolated colonies panels were inoculated according to CLSI recommendations (CLSI additional reference) and incubated in ambient air at 35±1°C for 16-20 h. Panel results were read visually, and minimal inhibitory concentrations (MIC) were determined.

#### MIC Value Processing

The bacterial culture may not grow for the smallest tested drug concentration (expressed as <= *x*) or show no significant growth decrease for the highest tested concentration (expressed as > *x*). In these cases, in order to allow consistent processing, the MIC values were transformed as follows: In case of <= *x*, the MIC value was set to *x* / 2 (e.g. “<=0.25” was set to “0.125”), and in case of > *x*, the MIC value was set to *x* * 2 (e.g. “>64” was set to “128”). Additionally, we considered only the MIC value of the first agent in case of drug combinations (e.g. “32/16” was set to “32”).

#### Drug Information

The 21 used drugs were grouped into 8 drug classes based on their category in the EUCAST guidelines[43]: 7 drugs belong to cephalosporins (cefazolin and cephalotin – 1st generation, cefuroxime – 2nd generation, cefotaxime, ceftazidime and ceftriaxone – 3rd generation, cefepime – 4th generation), 4 to penicillins, 3 to carbapenems, 2 to fluoroquinolones, 2 to aminoglycosides, 1 drug is a tetracycline, 1 is a monobactam and falls into category “miscellaneous” (**Supplementary Table S2**).

#### Resistance Classification

EUCAST guidelines[43] (v. 4.0) were used for MIC value classification. Isolates were classified as resistant, intermediate or susceptible. An isolate was considered to be resistant if the corresponding MIC value was greater than the resistance breakpoint. If the MIC value was below or equal to the susceptibility breakpoint the isolate was considered to be susceptible. If the MIC value was between the two breakpoints, the isolate was considered as “intermediate”. If no breakpoint was available for a specific drug and bacterial group, no classification was performed.

### Genome-based taxonomic classification

Kraken[44] (v. 0.10.4-beta) was used with the default database containing finished genomes from the NCBI RefSeq database (January 13^th^, 2015) and a k-mer length of 31. Report files were created from the raw output using “kraken-report” and processed to retrieve the following information: The first best species hit relative to the percentage of mapped sequences; the number of sequences mapped to best hit; the number of sequences classified at species level; the number of unclassified sequences; and the total number of reported sequences. In addition, sensitivity values, precision values, and percentages of unassigned sequences were calculated. Sensitivity was defined as the ratio of reads assigned to the best hit over the total number of reported reads. Precision was defined as the ratio of reads assigned to the best hit over reads classified at species level. For each sample, the taxonomic lineage from the species to the class level was retrieved using the R package “taxize”[45] and the NCBI[46] taxonomy database (accessed on February 8^th^, 2016). An overview of the taxonomic composition of the dataset was created using Krona[47].

### NGS processing and assembly pipeline

The raw reads were trimmed using Trimmomatic[48] (v. 0.35, command line parameters: PE ILLUMINACLIP:NexteraPE-PE.fa:1:50:30 LEADING:3 TRAILING:3 SLIDINGWINDOW:4:15 MINLEN:36). Trimmed paired-end reads were assembled *de novo* into scaffolds (from now on called contigs for simplicity) using SPAdes[49] (v. 3.6.2, parameters: -k 21,33,55 --careful) and annotated by Prokka[50] (v. 1.11, parameters: --gram neg --mincontiglength 200). Assembly quality was assessed using QUAST[51] (v. 3.2, parameters: --contig-thresholds 0,100,200,500,1000 --min-contig 200).

#### Mean assembly coverage

Trimmed reads were mapped to the contigs (minimal length 200 bps) using BWA[52] (v. 0.7.12) and SAMtools[53] (v. 1.2; command line: bwa mem –M –t <cores> <contigs> <forward reads> <reverse reads> | samtools view @ <cores> -bt <contigs> - | samtools sort -@ <cores> - <bam>), then coverage histogram was computed using BEDtools[54] (v. 2.25; parameters: bedtools genomecov –ibam <bam> -g <contigs>> <hist>). Finally mean coverage was computed over all contigs.

#### Essential genes

Essential genes as defined by Dupont *et al*.[33] were downloaded (https://github.com/MadsAlbertsen/multi-metagenome/raw/master/R.data.generation/essential.hmm, accessed March 7, 2017) and were searched in the present assemblies (protein FASTA files of translated CDS; *.faa) using hmmsearch from the HMMER software package (http://hmmer.org/, v. 3.1b2, parameters: --cut-tc). Only hits with at least one domain satisfying the reporting thresholds (column “rep” in table output files) were considered. Best hits for each isolate and essential gene were determined with respect to the reported full sequence e-value. Finally, each considered hit was assigned to a centroid — i.e., the centroid covering the gene from the corresponding hit.

#### Resistance factors

The Resfams core database[34] of HMMs (v1.2) was used to identify known resistance factors in the present assemblies (*.faa, FASTA file of protein annotations) using hmmsearch from the HMMER software package (http://hmmer.org/, v. 3.1b2, parameters: hmmsearch --cut_ga --tblout output.tblout Resfams.hmm input.faa > output.hmmout).

MLST. MLST profiles were determined using the BLASTn search-based tool mlst (https://github.com/tseemann/mlst, accessed August 8, 2016, v. 2.9, parameters: --minid 99 --mincov 75 --minscore 99) on assembled contigs (minimal length of 200 bps).

### Sample filtering

First, the bacterial isolate samples were filtered on the basis of their taxonomic assignment and assembly quality. For the taxonomic assignments, the minimal sensitivity was set to 50% (0% for *Shigella*), the minimal precision to 75% (60% for *Shigella*), and the minimal percentage of unclassified reads to 30%. The cutoff values were “relaxed” for *Shigella* because of the well-known problem of high genetic similarity between the *Shigella* species and *E*. *coli*[55] — i.e., it is difficult to differentiate between these organisms at the nucleotide level, which affects the taxonomic sensitivity. For the *de novo* assemblies, we used the criteria defined by RefSeq[37]: number of contigs ≤ 1,000, N50 ≥ 5,000, L50 ≤ 200. Samples that passed both filtering steps were grouped by their species taxon, and only species containing at least 50 samples were further considered, resulting in the following 18 species (referred to as “main species” in the manuscript) passing the filtering step: *Acinetobacter baumannii*, *Burkholderia cenocepacia*, *Citrobacter koseri*, *Enterobacter aerogenes*, *E. cloacae*, *Escherichia coli*, *Klebsiella oxytoca*, *K. pneumoniae*, *Morganella morganii*, *Proteus mirabilis*, *Pseudomonas aeruginosa*, *Salmonella enterica*, *Serratia marcescens*, *Shigella boydii*, *S. flexneri*, *S. sonnei*, *Staphylococcus aureus*, and *Stenotrophomonas maltophilia*. Additionally, samples containing more than 10 essential genes in multiple copies were examined more closely by running Kraken (k = 31) on the nucleotide sequences of the annotated genomic features (*.ffn). Report files were created from filtered assignments (kraken-filter, threshold 0.05) and inspected manually in order to determine whether a large percentage of sequences was assigned to unexpected species. In total, 8,729 isolates remained assigned to the 18 main species mentioned above.

### Pan-genome construction

Roary[17] (v. 3.5.7, parameters: -e -n -i 90 -cd 90 -a -g 70000 -r -s -t 11) was used to construct the species-level pan-genomes.

#### Centroid HMMs

The protein sequences were extracted from the FASTA files of translated CDS (*.faa) created by Prokka. For non-CDS sequences, protein sequences were created by translating the corresponding nucleotide sequence from the nucleotide FASTA files (*.ffn) using BioPython (parameters: table=11, stop_symbol=“*”, to_stop=False, cds=False). Multiple sequence alignments were created using MUSCLE[56] (v. 3.8.31, parameters: -maxiters 1 -diags -sv -distance1 kbit20_3), and HMM profiles were calculated using hmmbuild from the HMMER software package (http://hmmer.org/, v. 3.1b2).

### Database

The GEAR database was implemented using the Python Web framework Django (v. 1.9.5) and MySQL (v. 15.11) as the database management system. HMM search in Resfams core database and centroid HMM profiles is implemented using package/library HMMER (http://hmmer.org/, v. 3.1b1). Moreover, sketches of centroid nucleotide sequences were computed using Sourmash[35] (v. 2.0.0.a1, sketching parameters: sourmash compute --dna --singleton --scaled 10 --seed 42 --ksizes 21, indexing parameters: sourmash index --dna --ksize 21), and Mash/MinHash[38] (v. 1.1.1, default parameters) was used to create sketches of GEAR isolates, finished bacterial genomes from the NCBI RefSeq database (downloaded on June 17th, 2017 using the NCBI genome downloading scripts of Kai Blin (https://github.com/kblin/ncbi-genome-download, accessed October 18, 2017, v. 0.2.2, using “ncbi-genome-download --section refseq --assembly-level complete --human-readable --parallel 10 --retries 3 --verbose bacteria” with “--format fasta” and “--format cds-fasta”) including 7,118 bacterial genomes, and assembled bacterial genomes from the NCTC 3000 database of Public Health England and the Wellcome Trust Sanger Institute (downloaded on July 10^th^, 2017) including 1,052 bacterial genomes.

### Resistance profile analysis of cultured isolates from the Gram-negative collection

#### Drug correlations

Considering only species with at least 50 isolates, pairwise drug correlations were computed using the MIC value profiles (Spearman’s correlation coefficient, all samples and for each species taxon separately). Drugs with a single MIC value across all considered samples were removed prior to correlation computation. To visualize possible drug-drug associations, hierarchical clustering using Euclidean distance and average linkage was applied.

#### Association between isolate collection year and resistance profiles

Two-sided WMW-test (R package exactRankTests, v. 0.8-29) was applied to the isolates with assigned collection year belonging to a species taxon with at least 50 isolates (in total: 8,768 isolates from 18 taxa). The isolates were divided into resistant and non-resistant (susceptible or intermediate) groups. No test was performed if one of the groups included less than 10 isolates or all isolates in a group were collected in the same year. All *P* values were adjusted using FDR.

### Phylogenetic analysis

Essential genes, found in at least 99% of the isolates used to construct the pan-genomes, were identified. Protein sequences for the corresponding best hits were extracted for each essential gene and isolate. Multiple sequence alignments were computed using MUSCLE[56] (v. 3.8.31, parameters: -maxiters 1 -diags –sv -distance1 kbit20_3) for each essential gene separately and concatenated into one alignment. If an isolate did not have any matches, an empty alignment sequence (i.e., containing only gap characters) was added. RAxML[57] (v. 8.2.9, raxmlHPC-PTHREADS) was used to construct a phylogenetic tree from the aggregated alignment. After removal of sequence duplicates (2,297) and alignment columns containing only undetermined values (147), the tree was built using the CAT model (parameters: -p 12345 -m PROTCATAUTO -F -T 30).

### Pan-genome analysis

#### Centroid rate estimation

The centroid presence–absence tables created by Roary were used to estimate the median number of total, new, unique, and core centroids in species-level pan-genomes relative to the number of isolates used (rarefaction). For each pan-genome, the columns (isolates) of the table were permuted 100 times. Starting from the first isolate, centroid counts were calculated in a cumulative manner for each permutation. The following centroid categories were defined: total set of centroids, comprising centroids found in at least one of the included genomes; new centroids are centroids found only in the last included genome; unique centroids are centroids found only in one of the included genomes; core centroids are centroids found in at least 90%, 95%, and 99% of all included genomes. The median centroid counts were computed over all permutations. The curve of the total number of centroids was fitted using nonlinear least-squares estimates (R method “nls”) of the power law function *n= a N^Y^* (where *n* is the total number of centroids, *N* is the number of included genomes, and *a* and *y* are constants) to the median counts.

#### Two-dimensional embedding of pan-genome centroids

BusyBee Web[58] was used to represent the pan-genome centroids in two dimensions. In brief, pentanucleotide frequencies were computed and transformed into two dimensions using Barnes–Hut stochastic neighbor embedding[59]. Due to the use of centroids rather than contigs or long reads, the border points threshold and cluster points threshold were set to 500. Individual pan-genomes were mixed *in silico*, centroids with a frequency ≥ 90% were used as input to BusyBee Web, and the 2D coordinates were downloaded. Here, in addition to the sample frequency overlay, centroids were colored according to the respective species of the centroid’s source pan-genome.

### Resistance association analysis

#### Association between resistance profiles and centroid presence

All samples used to construct the pan-genomes and with a resistance profile were considered. Binary centroid presence/absence matrices were used as features. A species/drug combination was not analyzed if more than 90% of the samples were (not) resistant. The predictors were first filtered to remove (nearly) constant and correlated features and features with many missing values. All predictors with more than 95% missing values or with more than 95% of the entries having the same value (missing values ignored) were removed. Correlated features were removed by computing pairwise feature correlations (fastCor from R-package HiClimR, v. 1.2.3), clustering them by hierarchical clustering (distance = 1 – cor^2, average linkage), cutting the resulting tree at height 0.0975 (1 – (0.95^2)), and keeping only medoids (minimal average distance to other cluster members) within each obtained cluster. All features were scored using EIGENSTRAT[60] (v. 6.0.1) to correct for possible population structures. First, PCA was run to compute the top 50 principal components using only retained features. Then, the number of components (k) used for the subsequent computation was chosen such that the estimated genomic inflation factor (lambda) was below 1.1 for smallest possible k. If no of the computed lambda values was below 1.1 then k with the smallest lambda value was chosen. The value of k was successively increased from k=1 to k=50 by an increment of 2. With the chosen value of k, test statistics were generated for all features and *P* values were computed using the Chi-squared distribution with one degree of freedom. Finally, FDR adjustment was applied

#### Number of Resfams covered by the significant resistance association results

For each centroid with a significant resistance association result (adjusted *P* value below 1e-5), all hits from the centroid clusters members to the Resfams core database were retrieved. Subsequently, for each Resfam the number of unique centroids was counted including at least one cluster member with a hit to the corresponding Resfam.

### Application example

The assembly of the complete *K*. *pneumoniae* genome published by Kao et al.[39] (NCBI assembly ID ASM195283v1, RefSeq assembly accession GCF_001952835.1) was included in the collection of the finished bacterial genomes downloaded from the NCBI RefSeq database already described above. The genomic FASTA containing the chromosome and plasmid sequences was uploaded to the GEAR-base web-server for genome analysis using default parameters (https://gear-base.com/gear/pangenome/genomesearch/job=b568c458-f68a-4aa1-b78b-dad72dddfd5a/). The FASTA containing the nucleotide sequences of all CDS was uploaded for gene-based analysis with only Resfams search and Sourmash search in centroids enabled and using default parameters (https://gear-base.com/gear/pangenome/genesearch/job=0e42e149-a70d-4796-b40a-7f7168dc5077/). The nucleotide sequences of eight resistance genes reported by Kao et al. (WP_004176269.1, WP_076027158.1, WP_004146118.1, WP_000018329.1, WP_032491824.1, WP_000557454.1, WP_000976514.1, and WP_000027057.1) were saved in a separate FASTA file and uploaded for gene-based analysis with all options enabled and default parameters (https://gear-base.com/gear/pangenome/genesearch/job=d8792c0e-bbe7-4936-a7b7-c2846b727afe/).

### Availability

GEAR-base is freely available for academic research use after the user has registered and accepted the terms of use available under https://gear-base.com. Because of the sheer size and further legal and ethical constrains we cannot make all data fully accessible for batch download. If users should be interested in getting access to the sequencing raw data a special request in this respect is required. For this, we provide a respective request details on the GEAR-base homepage. The sequences of pan-genome centroids can be downloaded directly from the GEAR-base homepage. Custom scripts used for processing, analyzes and plotting can be found at https://github.com/VGalata/gear_base_scripts/.

## Authors’ contributions

VG performed the computational analysis, implemented the database, and drafted the manuscript together with CCL, who also contributed to the data analysis. GH-S and AF performed the next-generation sequencing of the isolates. MH and LvM. provided the *S*. *aureus* isolate collection. AEP, SS, CS, and AP provided the Gram-negative isolate collection. All of the authors reviewed the manuscript, provided comments, and approved the final manuscript.

## Acknowledgements

Research for this study was supported by Siemens Healthcare, the Curetis Group, and in parts by the Best Ageing Grant 306031 from the European Union as well as Austrian Research Promotion Agency Grants 866389 and 863729. We would like to thank Siemens Healthcare and the Curetis Group for their support and for the dataset provided. We are grateful to Laura Smoot, Andrea L. Mrotz, Khoa D. Nguyen, Michael A. Andora, Jose Enrique Fernandez, Nicholas E. Terzakis, Paula Swiatkowski, Usha Vajapey, and Stacie Ho for technical support. We also would like to thank Andy Ying and Gabriel Rensen for their support.

## Supplementary material

**Supplementary Figure S1 Local and temporal distribution of the isolates**

The numbers of isolates collected in (**A**) different countries and (**B**) years for species with least 50 isolates.

**Supplementary Figure S2 Drug correlations**

Heatmaps of the drug correlation matrices based on MIC value profiles. The first plot corresponds to correlation matrix computed using all isolates of selected species taxa; the remaining plots were generated for each species separately. Only species with at least 50 isolates were included. The cells in the heatmap are colored with respect to the correlation values (red for −1, blue for 1). The drugs were ordered using hierarchical clustering (Euclidean distance, average linkage) and colored with respect to their class.

**Supplementary Figure S3 Results of comparing temporal distribution of non-resistant and resistant isolates**

Two-sided WMW test comparing collection year distributions between non-resistant and resistant isolates. Only samples with collection dates available from species with at least 50 isolates were included. Heatmap of the negative log-transformed (base 10) adjusted (FDR) *P* values. The taxa and drugs were grouped by hierarchical clustering (Euclidean distance and average linkage on transformed *P* values). White cells contain non-significant adjusted *P* values (>=0.01) or omitted tests (e.g., due to insufficient group size). Cases where the test estimate was below zero were marked with “x”, otherwise – with “o”.

**Supplementary Figure S4 Temporal distribution of non-resistant and resistant isolates**

Boxplots of collection year for the non-resistant and resistant isolates for all species-drug combinations with a significant result. Only samples and drugs used for comparison using the two-sided WMW test are shown.

**Supplementary Figure S5 Frequency of essential genes per species**

Shown is the mean number of hits. Rows (species) and columns (essential genes) were ordered using hierarchical clustering (Euclidean distance, average linkage). The bar plot at the top of the heatmap shows the frequency of the essential genes over all isolates.

**Supplementary Figure S6 Phylogenetic tree of the isolates**

The phylogenetic tree of the isolates built using RAxML and multiple sequence alignments of the best hits of the essential genes found in at least 99% of the isolates. The tree tips are colored relative to the isolate species taxon.

**Supplementary Figure S7 Essential gene hits per sample and number of centroids covering essential genes**

(**A**) Essential gene hits per sample found in fewer than 99% of the isolates. The number of hits per isolate is shown in color and the isolates were grouped by species. The bar plot at the top of the heatmap shows the frequency of the essential genes over all isolates. (**B**) Mean number of centroids containing genes matched to essential genes per species. The bar plot shows the mean across all species. Bars above 1.25 are highlighted in red.

**Supplementary Figure S8 Frequency of Resfams hits per species**

Shown is the mean number of hits per resistance factor from the Resfams core database. The rows and columns were ordered using hierarchical clustering (Euclidean distance, average linkage). The bar plot at the top of the heatmap shows the frequency of the resistance factor across all isolates (frequencies of at least 90% are highlighted in red).

**Supplementary Figure S9 MLST analysis**

Pie charts showing the proportion of found MLSTs. The top 5 known STs are shown, known STs not included in the top 5 were grouped into “other”, “new” after the name of the scheme refers to new STs, and “unknown” refers to cases where no matching scheme was found.

**Supplementary Figure S10 Assembly length and pan-genome size**

The upper plot illustrates the median assembly length (solid black points), minimal and maximal assembly length (error bars), and pan-genome size (cumulative length over all centroids, diamond-shaped points) in Mb. The lower bar chart shows the number of centroids per pan-genome.

**Supplementary Figure S11 Estimated centroid rates for all pan-genomes**

The plots show the centroid rates of (**A**) all, (**B**) unique, (**C**) new, and (**D** – **F**) core (99%, 95%, 90%) centroids as a function of the number of genomes (isolates) included in the pan-genome. The rates were estimated from 100 permutations of the centroid presence–absence matrices as the median value over all permutations performed.

**Supplementary Figure S12 Two-dimensional embedding of per-species pan-genome centroids**

Centroids with a sample frequency >= 90% are shown. Colors represent the respective species of the centroids’ source pan-genome. Per-species medians are labeled.

**Supplementary Figure S13 Resistance associations and Resfams**

Number of centroids covering Resfams considering only centroids with a significant resistance association. **A**. Histogram of the centroid counts; the y-axis is square root transformed. **B**. The top 10 Resfams with their mechanism class and the number of centroids covering them.

**Supplementary Table S1 Taxonomy overview**

Taxonomy overview of the 11,087 isolates showing the Kraken-based species assignments and their lineages extracted from the NCBI taxonomy database.

**Supplementary Table S2 Drug overview**

Drug information table listing drugs included in the antibiotic susceptibility tests (AST) of the Gram-negative isolate collection. The table includes drug name, drug abbreviation, drug class, and the concentrations used in AST.

**Supplementary Table S3 Resistance data overview**

Overview of resistance data for the samples from the Gram-negative collection. The table includes only species with at least 50 isolates (prior to sample filtering). The columns are: Number of isolates, drugs with available resistance profiles (i.e. available MIC value breakpoints) and their number, number of susceptible isolates (i.e. excluding isolates without a resistance profile and with at least one resistance value), number of isolates resistant to one, two or at least 3 drugs. The values in brackets show the number of isolates after the sample filtering step; “NA” means that no isolates of the corresponding species were kept after the filtering step.

**Supplementary Table S4 WMW test results**

Statistics and results of the WMW tests of collection year distribution between non-resistant and resistant isolates. If no test was performed the corresponding cells are empty. The table includes group sizes, test statistics, estimates and confidence intervals, and raw and adjusted *P* values.

**Supplementary Table S5 Power law function estimation for the median centroid count**

Estimated power law functions using nonlinear least squares to describe the median number of all centroids per pan-genome as a function of the number of genomes included.

**Supplementary Table S6 Comparison of the number of centroids, and the number of samples between GEAR-base and panX**

The font color of species not found in panX was set to gray.

The columns are centroid count with frequency below 90%, at least 90% and total count in GEAR-base; number of samples in GEAR-base; centroid count with frequency below 90%, at least 90%, and total count in panX; number of strains in panX; the difference in centroid count between two resources for centroids with frequency of at least 90% and total centroid counts.

